# High Hedgehog signaling is transduced by a multikinase-dependent switch controling the apico-basal distribution of the GPCR Smoothened

**DOI:** 10.1101/2022.01.19.476759

**Authors:** Marina Gonçalves-Antunes, Matthieu Sanial, Vincent Contremoulins, Sandra Carvalho, Anne Plessis, et Isabelle Bécam

## Abstract

The oncogenic GPCR Smoothened is a key transducer of the Hedgehog morphogen, which plays essential roles in the patterning of epithelial structures. Here, we examine how Hedgehog controls Smoothened subcellular localization and activity in a polarized epithelium using the Drosophila wing imaginal disc as a model. We provide evidence that Hedgehog promotes the stabilization of Smoothened by switching its fate after endocytosis toward recycling. This effect involves the sequential and additive action of Protein Kinase A, Casein Kinase I and the Fused kinase. Moreover, in the presence of very high levels of Hedgehog, a second effect of Fused leads to the local enrichment of Smoothened in the most basal domain of the cell membrane. Together, these results link the morphogenetic effects of Hedgehog to the apico-basal distribution of Smoothened and provide a novel mechanism for regulation of a GPCR by plasma membrane subcompartimentalisation.

## INTRODUCTION

During development, signaling pathways control epithelial morphogenesis by acting on cell proliferation, differentiation, survival and migration. Epithelial cells uniquely display an apico-basal (Ap-Ba) polarity with differential distribution of phospholipids and protein complexes between the different membrane domains (for review see Ikenouchi, 2018). This leads to functionally separated subregions with distinct properties and physiological functions, such as the microvilli in the apical domain, cell-cell adhesion junctions in the lateral domain, and cell-matrix adhesion in the basal domain. Finally, the apical and basal regions are in contact with extracellular environments, which can differ in the nature and dose of signaling molecules. For all these reasons, the control of signaling receptor distribution among these specific subdomains of the plasma membrane is expected to be critical for correct signal transduction.

The conserved Hedgehog (HH) signals play major roles in the development of metazoans. Initially identified in the fly model, HH signaling is involved in the promotion, development and/or metastasis of numerous types of tumors and drugging this pathway is a major goal for cancer therapies (for review see Briscoe and Thérond, 2013). In flies, HH controls the patterning of many structures including the wing imaginal disc (WID), which has been instrumental for the study of HH signaling (for review see Hartl and Scott, 2014). In this epithelial structure, HH emanating from the posterior (P) cells signals to the anterior (A) cells near the A/P boundary, thus controlling the expression of target genes in a dose-dependent manner. Of note, HH molecules form two gradients in the WID: an apical gradient, required for long-range, “Low HH” responses that depend on glypicans, and a basal one, for short-distance, “High HH”responses, that is based on the transport of HH by exosomes associated to filopodia-like structures oriented in the A/P axis (D’Angelo et al., 2015; González-Méndez et al., 2017).

HH transduction requires Smoothened (SMO), a G Protein-Coupled Receptor (GPCR) like protein. In the absence of HH, the HH co-receptor Patched (PTC) inhibits SMO, probably by depleting accessible cholesterol from the outer leaflet of the plasma membrane (Kinnebrew et al., 2021 and see for review Radhakrishnan et al., 2020), which promotes the formation of a repressor form of the transcription factor Cubitus interruptus (CI). The binding of HH to PTC inhibits its negative effect, which blocks the cleavage of CI, leading to the transcription of pathway target genes by full-length CI. SMO acts as a scaffold to transduce HH signaling to CI via an intracellular complex (called HTC for Hedgehog Transduction Complex) bound to its cytoplasmic C-terminal tail (cytotail), and which includes CI and a protein kinase called Fused (FU) (Malpel et al., 2007; Robbins et al., 1997; Sisson et al., 1997). SMO activation is associated with conformational switches, both of its cytotail and of its extracellular domains; these events correlate with its clustering, which seems critical for the downstream activation of the pathway (Fan et al., 2012; Shi et al., 2011; Su et al., 2011; Zhao et al., 2007).

Several labs -including ours-have highlighted the role of endocytic trafficking and the importance of post-translational modifications in the regulation of SMO’s levels, localization and activation. SMO activation in the presence of HH is associated with changes in its localization: from internal vesicles to the plasma membrane in *Drosophila* and from the cell body to the primary cilium in mammals (Denef et al., 2000; Huangfu et al., 2003). Both events are controlled by extensive phosphorylation of SMO’s intracellular tail by multiple kinases (for review see Chen and Jiang, 2013). Despite significant differences, the processes involved are remarkably conserved as illustrated by the fact that human SMO (hSMO) can be relocalized in response to HH to the surface of fly cells (De Rivoyre et al., 2006). In Drosophila, many kinases (Protein Kinase A (PKA), Casein Kinase I (CKI), G-protein-coupled receptor kinase 2, Casein Kinase 2, Gilgamesh, atypical Protein Kinase C (aPKC)) are implicated, which regulate SMO activation and accumulation at the membrane (Apionishev et al., 2005; Chen et al., 2010; Jia et al., 2010, 2004; Li et al., 2016; Maier et al., 2014; Zhang et al., 2004). We have also identified a phosphorylation-based positive feedback loop between SMO and the FU kinase, which is required for the response to the highest doses of HH (Alves et al., 1998). In this process, an initial activation of SMO promotes the recruitment at the cell membrane of FU and its activation, which then further phosphorylates SMO, leading to an enhanced accumulation of the SMO/FU complex at the cell surface and high signaling activation (Claret et al., 2007; Sanial et al., 2017).

Are these events polarized along the apico-basal axis and what is their link with the gradients of HH? Previous studies indicated that SMO is unevenly distributed along the apico-basal axis of the wing imaginal disc epithelial cells (Denef et al., 2000; Jiang et al., 2014; Sanial et al., 2017). Here, by specifically labeling the population of SMO at the plasma membrane, we show that this uneven distribution is due to the cell surface population of SMO with its “high HH”-dependent accumulation in the most basal region. Moreover, blocking the endocytosis of SMO or following its fate after endocytosis reveals that SMO is initially targeted to the apical membrane. We show that HH does not dramatically affect SMO endocytosis but affects its post-endocytic fate, favoring recycling over degradation. Our results also indicate that in presence of high HH, a fraction of SMO accumulates in the most basal region. This and the analysis of various phosphomutants of SMO support a model whereby (i) HH enhances the recycling of endocytosed SMO in a phosphorylation level-dependent manner and (ii) very high levels of HH lead to FU-dependent basal trapping of SMO.

## RESULTS

### High levels of HH promotes a basolateral enrichment of cell surface SMO

To analyze the apico-basal subcellular localization of SMO, we used a *SNAP-smo* transgene, which encodes a fusion between the extracellular N-terminus of SMO and the enzymatic self-labeling SNAP-tag (Tirat et al., 2006). This SNAP-SMO fusion was previously shown to be functional *in vivo* (Sanial et al., 2017). Using different SNAP ligands allows one to label the whole SNAP-SMO population (hereafter called Total SNAP-SMO), or only the fraction of SNAP-SMO at the cell surface (Surf SNAP-SMO), or both Surf SNAP-SMO and the intracellular SNAP-eSMO (Intra SNAP-SMO) differentially (Figure S1) (Sanial et al., 2017). After imaging XZ sections of the labeled wing imaginal discs, we quantified SNAP-SMO mean intensity (sum of pixel values over the number of pixels) and integrated intensity (sum of pixel values) in three regions along the apico-basal axis: (i) the apical region (estimated here as the 15% most apical region), (ii) the basal region (arbitrarily defined as being the 10 % most basal part) and (iii) the lateral region in between the two other regions (see Figure S1A). Moreover, we systematically compared cells that do not receive HH in the far anterior region (FA) to the HH-producing posterior (P) cells that are used as a model for HH responsive cells, although they do not express HH target genes, due to lack of *ci* expression (Méthot and Basler, 1999).

First, we labeled Total SNAP-SMO. As shown in Figure 1 (1A, 1A”, 1B, 1B’, 1B”), it is not homogeneously distributed along the apico-basal axis, with a stronger apical accumulation in all the cells of the wing pouch and a increase in the basal region of the anterior in presence of high levels of HH. As expected, it accumulates in the posterior cells where it is 1.5 times more abundant than in the far anterior cells (Figures 1A, 1A”, 1B, 1B’, 1B” and 1E). The increase in the posterior region includes all the subregions along the apico-basal axis but was less pronounced in the apical region than in the lateral and basal regions (with 1.3, 1.5 and 1.6 fold increases, respectively) (Figure S1D). The relative intensities (calculated as the ratio of the integrated density of each region along the apico-basal axis over the integraded intensity of the three regions together, called column) show also that the fraction of apical SNAP-SMO is lower in the posterior region than in the far anterior region (Figure 1E’). This effect is associated with a slight (but not significant) increase in the relative abundance of the basal and lateral populations in the posterior cells compared to the far anterior one. Notably, these differences in the apico-basal distribution in the far anterior and posterior regions are confirmed when looking, by immunolabeling, at endogenous SMO (Figures S1B, S1C and S1C’).

**Figure 1:**
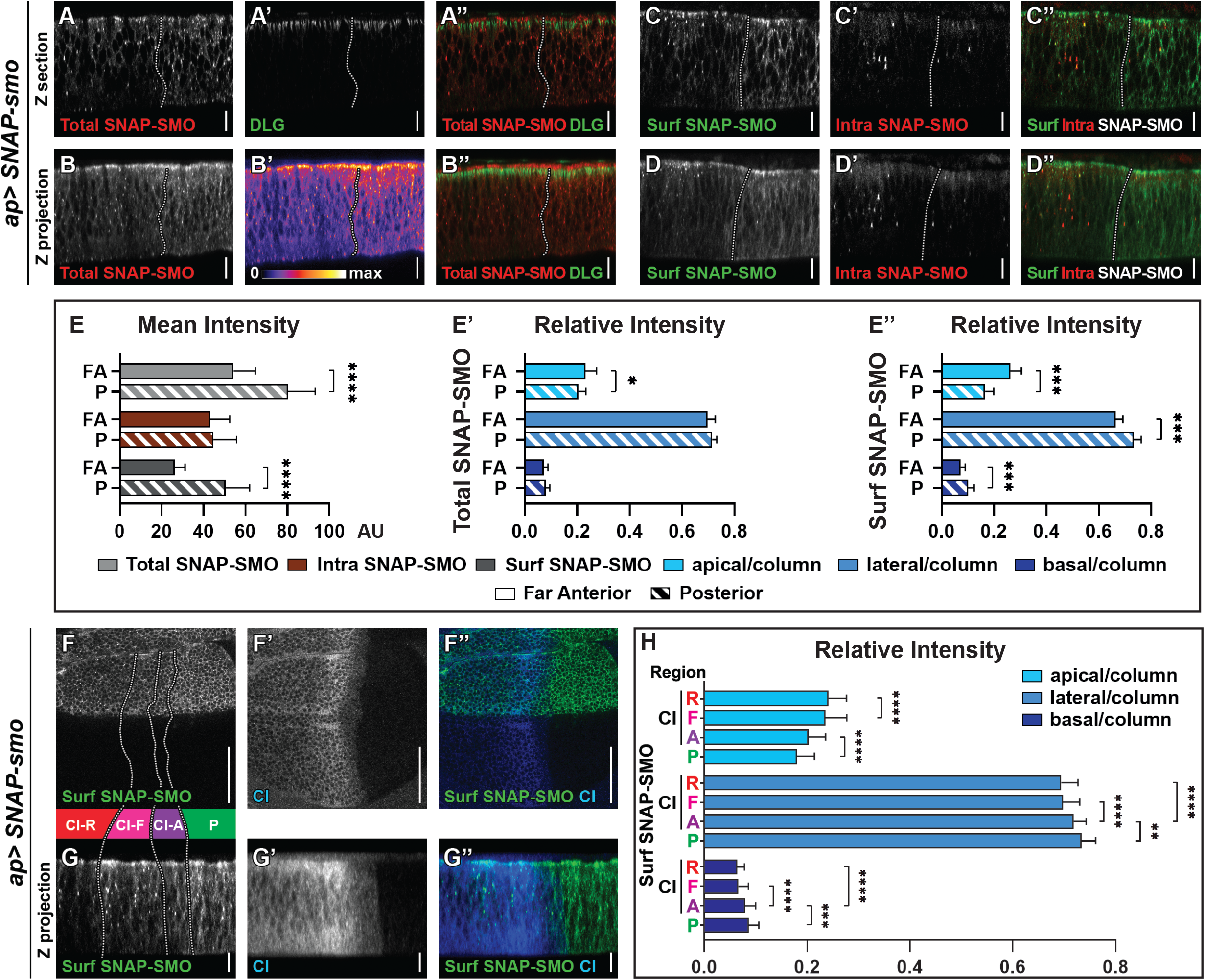
Cell surface SMO is unevenly distributed along the apico-basal axis and high levels of HH promotes its basolateral enrichment. **(A-B”)** Confocal images of third instar larva imaginal wing discs expressing *UAS SNAP-smo*^*wt*^ (driven by *apGal4)* in the dorsal compartment and labeled with a SNAP cell-permeable fluorescent substrate (A, B and red in the merge images A”, B”) and DLG (A’ and green in A’’ and B’’). (B’) Total SNAP-SMO in false colour. (A-A”): XZ section, (B-B”): XZ projection. **(C-D’’)** Confocal images of imaginal discs expressing *UAS SNAP-smo*^*wt*^ and labeled with a non-liposoluble fluorescent SNAP substrate followed by a cell-permeable fluorescent SNAP substrate. C-C” XZ section, D-D” XZ projection. Here and in the Figures 2 to 4: XZ sections are antero-posterior Z sections in the dorsal compartment. The disc images are oriented anterior to the left and apical up. The dotted vertical line represents the A/P boundary (determined by CI staining). The Z projection images correspond to an average intensity projection of 8 sections. The scale bars represent 50 μm for XY images and 10 μm for XZ images. **(E-E”)** Quantification of SNAP-SMO localization in the XZ sections. (E) Density (mean intensity) of total (pale grey), intracellular (Intra, red) and surface (Surf, dark grey) SNAP-SMO in two different regions of the wing imaginal disc: far anterior (FA) region (empty bars here and in Figures 2-4) where no HH is present and posterior (P) region (bars with diagonal lines here and in Figures 2-4. (E’ and E”) Distribution (intensity reported to the intensity of the whole column taken for quantification) of Total SNAP-SMO (E’) or of surface SNAP-SMO (E’’) in the apical (light blue), lateral (medium blue) and basal (dark blue) domains in the FA or P regions. Total SNAP-SMO: N=9 discs, for Intra and Surf: N=12 discs. The error bars represent the S.D. and the statistical analysis was performed using paired t-test for the mean intensities and Wilcoxon matched-pairs signed rank tests for the relative intensities. **(F-F”)** Confocal images of imaginal discs expressing *UAS SNAP-smo*^*WT*^, labeled with a non-liposoluble fluorescent SNAP substrate (F, G green in the merge images F”and G”) and immunolabeled for unprocessed CI (F’, G’, blue in F”and G”). XY images in (F to F’’) with anterior left and dorsal up. XZ images in (G to G”).The disc is divided in four regions (separated by dotted lines) along the anterior posterior axis based on CI staining: the posterior (green), and three anterior regions: CI-A (low levels of full-length activated CI, in purple), CI-F (higher levels of CI full-length, in pink) and CI-R (CI repressor not detected by the anti CI antibody used here, in red). See also Figure S1E. **(H)** Graph showing the relative intensity of surface SNAP-SMO in the apical, lateral and basal domains in CI-R, CI-FL, CI-A and P regions. N= 33. For all Figures, the precise genotypes are given in the Table S1, the statistical tests used and the p-values are indicated in the Table S2. See also Figure S1.

As HH promotes the accumulation of SMO at the cell surface, we set out to determine (i) whether the Surf SNAP-SMO and Intra SNAP-SMO populations are differentially distributed along the apico-basal axis and (ii) whether the apico-basal distribution of one or both is affected by HH. For that purpose, we labeled Surf SNAP-SMO and Intra SNAP-SMO with different fluorochromes (Figures S1F-F”‘). Intra SNAP-SMO is present in vesicle-like dots that are mainly in the lateral region of the cells (Figures 1C’, 1C”, 1D’, 1D”). Its abundance and localization are comparable in the posterior and far anterior regions (Figures 1E and S1D’). By contrast, the abundance, subcellular localization and apico-basal distribution of Surf SNAP-SMO are all different in the posterior and far anterior regions (Figures 1C, 1C”, 1D, 1D”, 1E, 1E”). In the far anterior region, it is present in dot-like structures at or near the apical region. These structures may correspond to Surf SNAP-SMO molecules endocytosed during the labeling. In the posterior region and in the anterior cells abutting the A/P boundary, Surf SNAP-SMO is more abundant than in far anterior (1.9 fold increase), mostly along the cell membrane, with fewer vesicle-like structures. Here also, a large fraction of Surf SNAP-SMO is at the apical membrane (see the quantification in Figure S1D”). However, despite this and the fact that the posterior increase in Surf SNAP-SMO occurs all along the apico-basal axis, this increase is stronger in the lateral and basal regions than in the apical one (fold increases: apical 1.2, medial 2.1, basal 2.6), leading to a relative decrease in the apical fraction and a concomitant relative enrichement of lateral and basal fractions. Since the intracellular population of SMO is not affected by HH, we focused the rest of our work exclusively on Surf SNAP-SMO.

Next, since HH acts as a morphogen in the wing disc, we tested whether HH also acts on the apico-basal distribution of Surf SNAP-SMO in a dose-dependent manner. For that, we quantified Surf SNAP-SMO in four regions across the wing disc epithelium, based on the immunodetection of the transcription factor CI (Figures 1F, 1G and S1A): the posterior compartment (CI not expressed) and three anterior regions: CI-R (corresponding to the far anterior region) where CI is processed into its shorter repressor form that is not detectable with the antibody used here), CI-F, corresponding to cells that receive medium-low levels of HH that lead to the stabilization of full-length CI and CI-A that corresponds to a full-length activated form of CI which is however, present at low levels. Both the relative and mean intensities of Surf SNAP-SMO distribution in these four regions show that HH acts in a dose-dependent manner on the apico-basal distribution of Surf SNAP-SMO (Figures 1H and S1E). Of note, this further validates our use of the posterior compartment to study the effects of HH on the apico-basal localization of SMO.

In summary, these data provide evidence that SMO is asymmetrically distributed along the apico-basal axis with higher levels in the apical region, and that HH increases the basolateral localization of surface SMO.

### HH controls the fate of SMO post-endocytocis

To understand how the distribution of Surf SNAP-SMO along the apico-basal axis is established, we looked at the consequences of blocking its endocytosis.

For that purpose, we first used a thermo-sensitive mutation of *shibire (shi*^*ts*^), a Drosophila Dynamin orthologue, which is central for the scission of coated vesicles (van der Bliek and Meyerowrtz, 1991). Blocking SHI activity -for less than an hour-leads to an accumulation of Surf SNAP-SMO in both compartments of the wing imaginal disc, with a stronger accumulation in the apical region of the cells (Figures 2A, 2B, S2A and S2B). Quantification of the mean intensities reveals that the increase in Surf SNAP-SMO levels is comparable in the posterior and far anterior regions. In both regions, this increase occurs all along the apico-basal axis of the cells (Figure 2C), with however a stronger increase in the apical region (with a 1.6 fold increase in both the far anterior and posterior regions, compared to the *shi*^*+*^ control at the restrictive temperature), than in the lateral (1.2 fold increase, both in far anterior and posterior regions) and basal regions (1.4 fold and 1.2 fold increases in the far anterior and posterior regions, respectively). This leads to a relative enrichment of Surf SNAP-SMO in the apical region of the cells, which is associated with its relative decrease in the lateral region (see the relative intensities in Figure 2C’). Note that the effects seen here are specific to the inactivation of *shi*^*ts*^ as no difference is observed at 18°C (Figure S2E).

**Figure 2:**
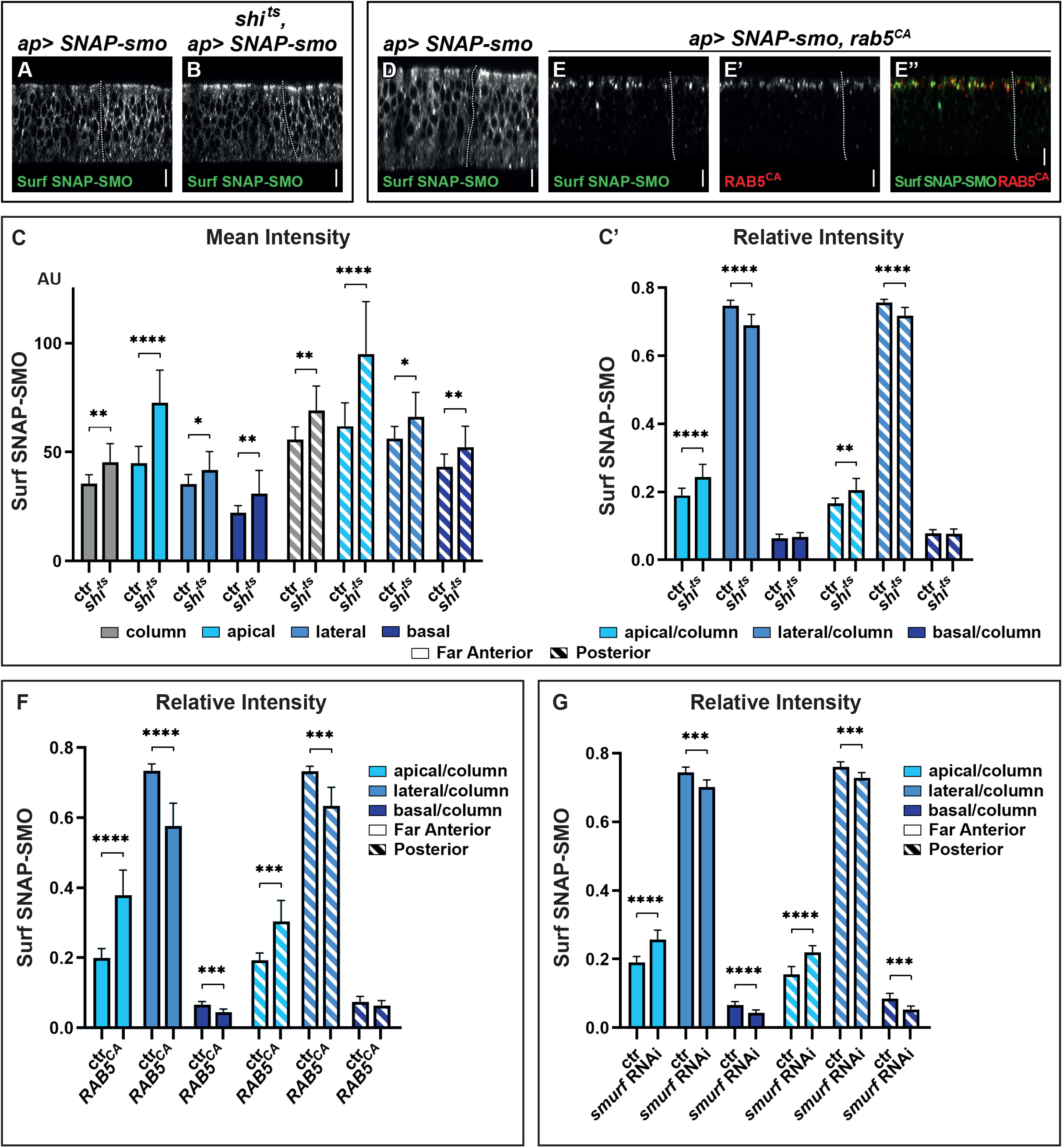
Blocking SMO endocytosis favors its accumulation in the apical region independently of HH. **(A-B)** XZ confocal images of wing imaginal discs from *shi*^*+*^; *apGal4; UAS SNAP-smo*^*WT*^ (A, called *ctr* for control in C and C’) or *shi*^*ts*^; *apGal4; UAS SNAP-smo*^*WT*^ (B, called *shi*^*ts*^ in C and C’) male flies put at restrictive temperature before dissection, labeling of surface SNAP-SMO and imaging under the same conditions. **(C-C’)** Quantification of the mean intensities (C) and relative intensities (C’) of Surf SNAP-SMO in the different regions of the disc (FA and P) and along the apico-basal axis, as indicated. The mean intensity is also shown for the whole epithelial column. N= 17 and 10 for *shi*^*+*^ and *shi*^*ts*^ flies, respectively. Note the weaker basal effect of SHI inactivation in presence of HH, which indicates that HH may slightly reduce SMO basal endocytosis. **(D-E”)** XZ confocal images of wing imaginal discs from *apGal4, Gal80*^*ts*^; *UAS SNAP-smo*^*WT*^ (D) or *apGal4, Gal80*^*ts*^*/UAS YFP-rab5*^*CA*^; UAS *SNAP-smo*^*WT*^ (E-E”) flies put at restrictive temperature (to allow *rab5*^*CA*^ expression) before dissection and labeled for surface SNAP-SMO. In the latter case, E” is a merge image with Surf SNAP-SMO in green and YFP-RAB5^CA^ in red. Note that for a better visualization of SMO, imaging of rab5^CA^ and control discs were not done in the same conditions. **(F)** Relative intensity of Surf SNAP-SMO along the apico-basal axis in *apGal4, Gal80*^*ts*^; *UAS SNAP-smo*^*WT*^ called ctr (n=11) and *apGal4, Gal80*^*ts*^*/UAS YFP-rab5*^*CA*^; UAS *SNAP-smo*^*WT*^ called Rab5^CA^ (n=8). **(G)** Relative intensity of Surf SNAP-SMO along the apico-basal in *apGal4; UAS SNAP-smo*^*WT*^ called ctr (n=20) and *apGal4/ UAS smurf RNAi; UAS SNAP-smo* ^*WT*^ called *smurf* RNAi (n=6) flies. Note that *Rab5*^*CA*^ or RNAi *smurf* overexpression have a much stronger effect than *shi*^*ts*^ inactivation, reflecting that endocytosis is blocked for a much shorter time in the latter case. Here and in all the following figures, the XZ images correspond to a unique XZ section. See also Figure S2.

To confirm these results, we also blocked endocytosis using a constitutively active Rab5 that is locked in the GTP bound state and induces endocytosis defects (D’Angelo et al., 2015). When endocytosis is blocked for 24 hours, Surf SNAP-SMO strongly accumulates in Rab5^CA^ positive endocytic vesicles, which are almost exclusively located in the apical region (Figures 2D, 2E-E”, S2C and S2D-D”). This leads to an decrease in the relative abundance of its lateral fraction and to a lesser extent, of its basal fraction (Figure 2F). These effects are seen both in presence (P region) and absence of HH (FA region) but are slightly weaker in its presence.

Finally, we ruled out an indirect effect of a general block of endocytosis, as we obtained similar results when we specifically blocked the endocytosis of SNAP-SMO by downregulating (by RNA interference, RNAi) the expression of *smurf*, which encodes a E3 ubiquitin ligase known to promote SMO endocytosis by mediating its ubiquitylation (Li et al., 2018) (Figure 2G).

In conclusion, these data reveal that (i) SMO endocytosis is not dramatically affected by HH and (ii) SMO endocytosis occurs all along the apico-basal axis but more apically than basolaterally. They also show that newly synthesized SMO is intially addressed and subsequently primarily endocytosed at the apical membrane. Moreover, the increase of the apical fraction of SMO due to blocking endocytosis and its associated reduction in the lateral and basal fractions, strongly suggest that a significant fraction of the apically endocytosed SMO is transcytosed to the basolateral membrane.

### Endocytosed SMO is targeted from the apical to the basolateral region

The above data indicate that the strong stabilization of SMO in response to HH is likely due to a reduction of its degradation after endocytosis. To study the fate of Surf SNAP-SMO after its endocytosis, we performed an endocytosis assay in which Surf SNAP-SMO labeling was followed by a chase. After the chase, the subcellular localization of Surf labeled SNAP-SMO shifted: its presence at the cell surface decreases and this is associated with increased localization in dot-like structures that likely correspond to endocytic vesicles (Figures 3A and 3B). The global levels of Surf labeled SNAP-SMO also decrease during the chase in the whole disc with a reduction of 46% in the far anterior cells (no HH) but of only 25% in the posterior cells (with HH) (Figure 3C). In the absence of HH, the apical region of the cells is slightly more affected than the lateral and basal regions (48% decrease in the apical region compared to 45% and 36% in the lateral and basal regions, respectively). This leads to an increase of the relative abundance of Surf labeled SNAP-SMO in the basal region (associated to a slight but non-significant decrease of the apical fraction). By contrast, in the presence of HH (P region), the reduction is much more pronounced in the apical region (40% decrease) than in the lateral (24% decrease) and basal region (which is barely affected, 8% decrease) (Figure 3C). This leads to a relative redistribution in the P region (compare to the FA region) of endocytosed Surf labeled SNAP-SMO from the apical to the basolateral region (Figure 3D).

**Figure 3:**
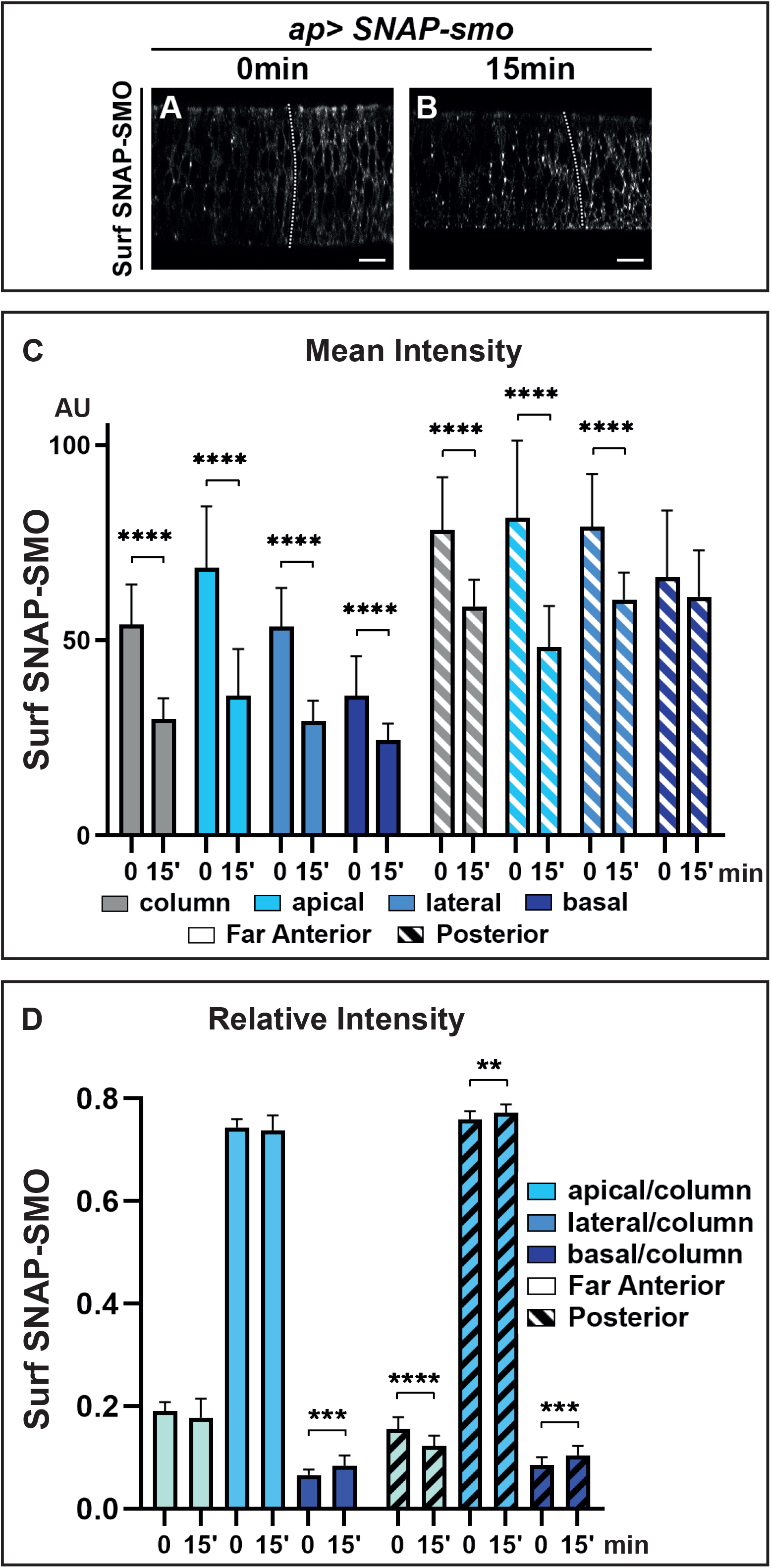
HH controls the fate of endocytosed SMO, favoring its recycling and leading to its basolateral accumulation. XZ confocal images of dissected wing imaginal discs expressing *UAS SNAP-smo*^*WT*^ *fixed* immediately (0 min) **(A)** or 15 minutes **(B)** after labeling of Surf SNAP-SMO. Quantification of the mean intensity **(C)** and of the relative intensity **(D)** in the far anterior and posterior regions of the disc and along the apico-basal axis. N= 20 for 0 min and 23 for 15 min.

In summary, our data show that, both with and without HH, SMO endocytosis leads to a change in its distribution along the apico-basal axis, in favor of the basolateral region, further suggesting that it is transcytosed. Moreover, HH reduces the degradation of endocytosed SMO and favors its presence in the lateral and basal regions. As HH does not dramatically affect SMO endocytosis, this indicates that HH controls the fate of endocytosed SMO by reducing its degradation and increasing its recycling.

### Phosphorylation by the PKA/CKI and FU kinases regulates the apico-basal localization of SMO

Given the importance of the phosphorylations of the cytotail of SMO, we analyzed the effect of these phosphorylations on the apico-basal localization of SMO by looking at forms of SMO mimicking (S to D replacements) or blocking (S to A substitution) these phosphorylations.

First, we looked at the constitutively active SNAP-SMO^PKA-SD^, which mimics SMO fully phosphorylated by PKA and CKI kinases (Jia et al., 2004). As expected from previous data with SMO^PKA-SD^ in cultured cells (Jia et al., 2004; Sanial et al., 2017), Surf SNAP-SMO^PKA-SD^ accumulates at the cell membrane both in the anterior and posterior compartments of the wing disc (Figures 4A, 4B, S3A-A’ and S3B-B’). Moreover, even in the absence of HH (FA cells), its accumulation in the apical region decreases in favor of the lateral and even more of the basal region, similarly to what is seen with Surf SNAP-SMO^WT^ in presence of HH (Figures 4E-E”). These effects in the far anterior region are partially suppressed by mutations that prevent the phosphorylation by the FU kinase, with especially a strong reduction of the basal localization of Surf SNAP-SMO^PKA-SD FU-SA^, compared with Surf SNAP-SMO^PKA-SD^ (Figures 4C, 4E-E” and S3C-C’). In contrast, Surf SNAP-SMO^PKA-SD FU-SD^, which accumulates at high levels at the cell surface in both compartments of the discs, shows a further decrease in its accumulation at the apical and lateral cell surface in anterior cells compared to SMO ^PKA-SD^ (Figure 4D, 4E-E” and S3D-D’). This effect is associated with a much strong basal enrichment.

**Figure 4:**
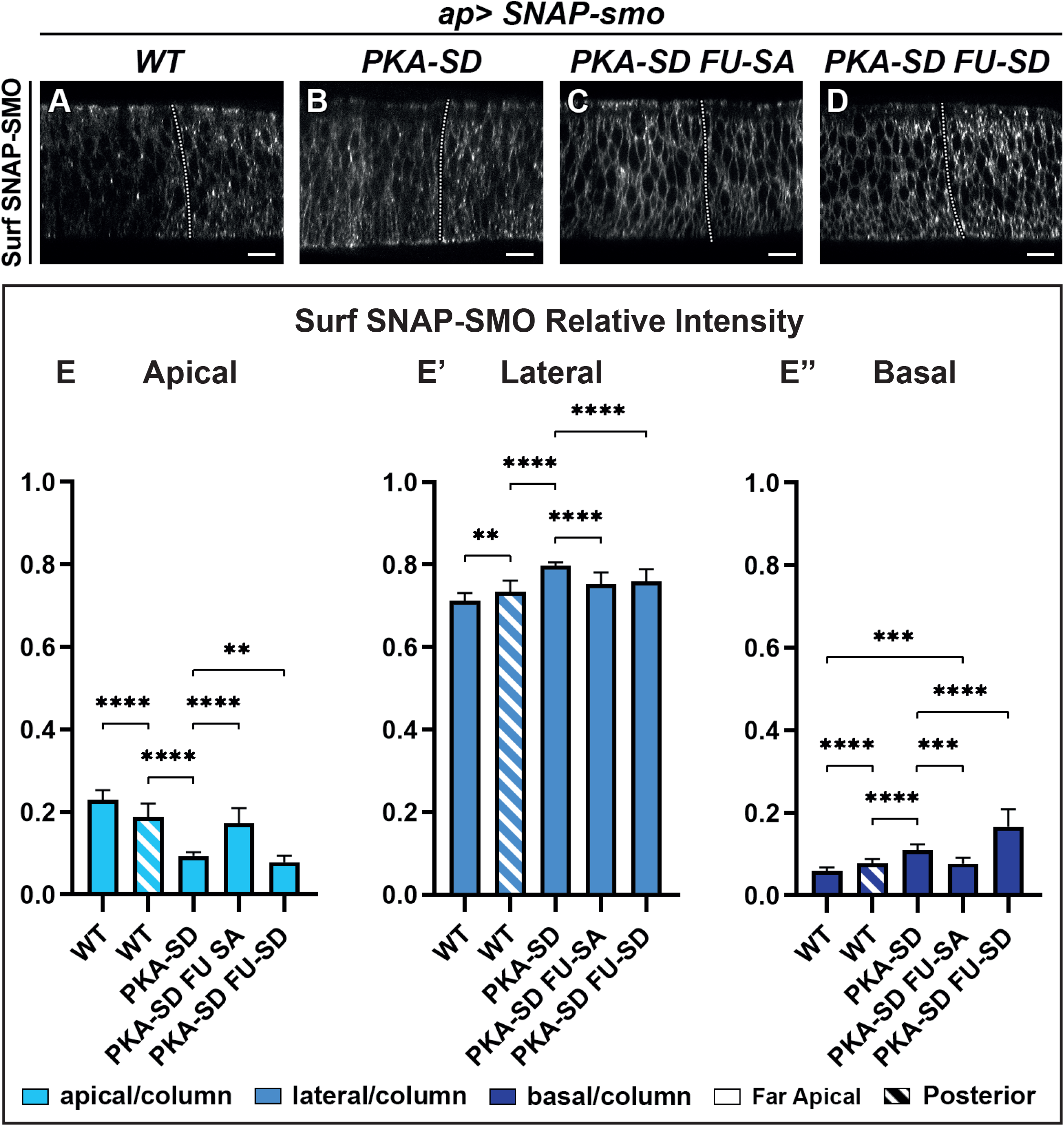
The apico-basal distribution of SMO is controlled by phosphorylation by PKA/CKI and FU kinases. **(A-D)** Confocal images of wing imaginal discs labeled for Surf SNAP-SMO^WT^ (A), SNAP-SMO^PKA-SD^ (B), SNAP-SMO^PKA-SD FU-SA^ (C), SNAP-SMO^PKA-SD FU-SD^ (D). Note that for a better visualization of SMO, imaging of the different forms of SNAP-SMO were not done under the same conditions. **(E-E”)** Quantification of the apico-basal distribution of the different forms of SNAP-SMO, as indicated. N=21, 10, 15 and 18 discs respectively. See also Figure S3. Note that the effects of the *PKA-SD* mutations are even stronger than the effect of HH on SNAP-SMO^WT^, probably due to the fact that in presence of HH, only a fraction of the SMO population is phosphorylated (Sanial et al., 2017).

Overall these results provide evidence that the phosphorylation by the PKA/CKI recapitulates the effects of HH on Surf SMO apico-basal distribution, leading to a relative apical depletion and basolateral enrichment. They also show that this effect is dependent upon further phosphorylation of SMO by FU, which especially enhances the localization of SMO in the basal region.

### FU is required first apically and then basally to promote high SMO activity

The FU kinase is present everywhere along the apico-basal axis, with some enrichment in the apical region. It is both diffused in the cytoplasm and present in vesicular puncta, some of which colocalize with SMO (Claret et al., 2007) (Figure S4A-A” and S4B-B”). To understand where FU acts on SMO, we trapped it in the apical or in the basolateral region of the cells using the GRAB bipartite system (Harmansa et al., 2017). In this method, GFP tagged FU (GFP-FU, known to be fully functional (Malpel et al., 2007)) is trapped in either the apical or basolateral domain via its binding to an intracellular GFP nanobody fused to an apical (T48) or basolateral (NVR1) transmembrane domain. For that purpose, we expressed both FU-GFP and its trap in the dorsal part of the disc and we immunolabeled endogenous SMO (Figure 5, note that in B-B”‘ and D-D”‘, the Z sections correspond to anterior YZ sections along the dorso-ventral axis, with the ventral compartment serving as an internal control).

**Figure 5:**
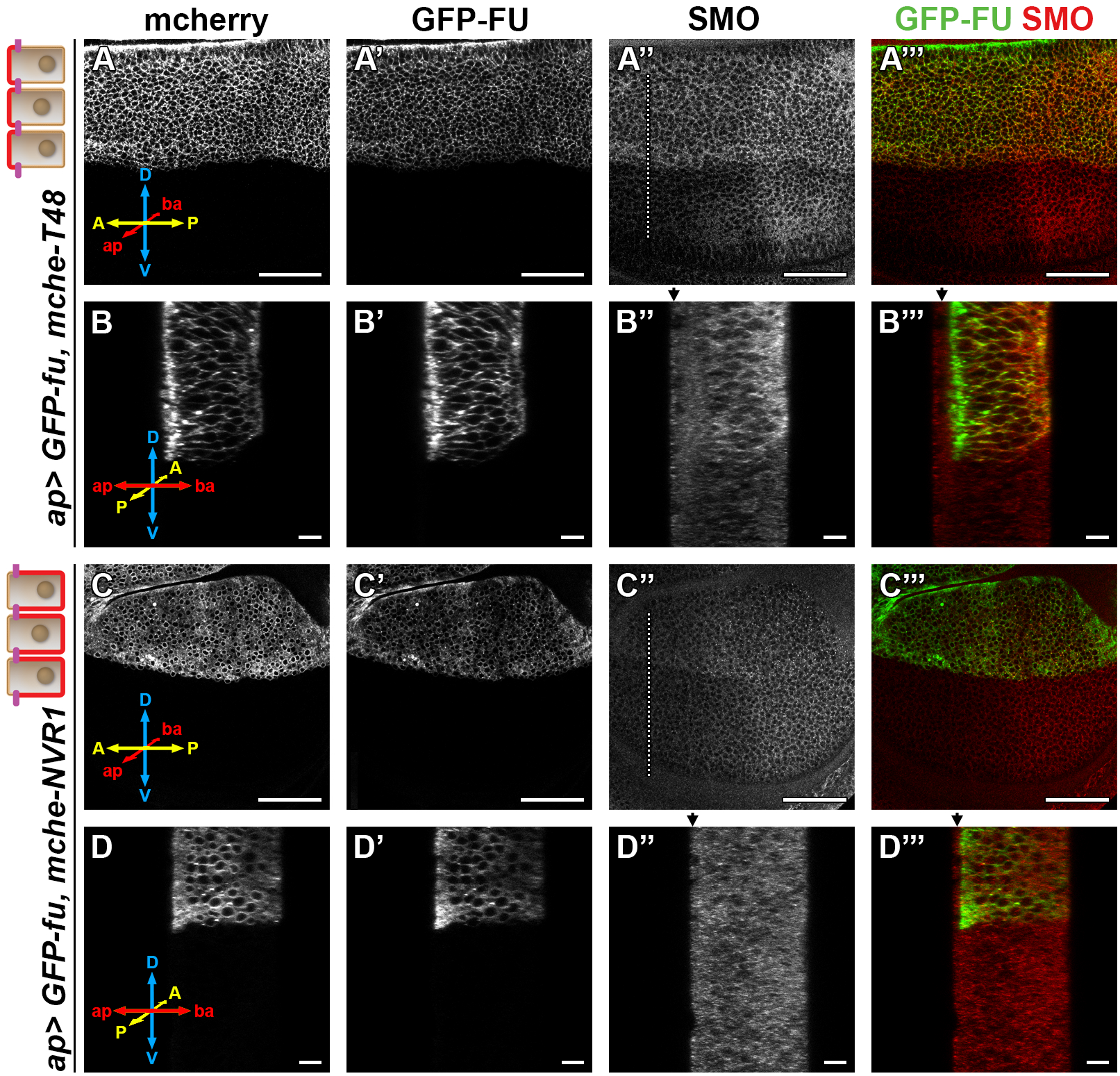
Trapping the FU kinase to the apical region promotes the stabilisation of SMO at the cell surface. Confocal images of wing imaginal discs coexpressing (using the apGal4 driver) *GFP-fu* and *T48* or *NVR1* fused to the *mcherr*y (*mche*). XY sections are shown in **(A-A”‘ and C-C”‘)**, anterior YZ section in **(B-B”‘and D-D”‘)**. mcherry is shown in (A, B, C and D), GFP-FU in (A’, B’ C’ and D’, green in the merge images A”‘, B”‘, C”‘ and D”‘) and immunolabeled endogenous SMO in (A”, B”, C” and D”, red in the merge images). Note that contrary to SNAP-SMO, endogenous SMO is also present in the peripodial membrane (arrow). D: dorsal, V ventral, A: anterior, P: posterior, ap:apical and ba: basal. See also Figure S4.

As expected, expression of GFP-FU with T48 leads to a strong apical localization of GFP-FU (Figure 5B’ and S4C’) while its coexpression with NVR1 leads to its basolateral enrichment (Figure 5D’ and S4D’). Apical tethering of GFP-FU (with T48) promotes the accumulation of SMO both in the anterior (Figure 5A”, 5B” and S4E) and posterior (Figure 5A”, S4C’” and S4E’) compartments. This effect is homogeneous along the apico-basal axis, with only a slight increase of the basal relative distribution of SMO (Figure S4E”). By contrast, basolateral trapping of GFP-FU (with NVR1) has only a slight effect on SMO accumulation (Figure 5C”-D” and S4D”‘) and seems to induce its vesicular localization.

Next, we tested the effects of trapping GFP-FU with T48 or NVR1 on HH signaling. We monitored the accumulation of CI and the expression of the low-HH target *dpp* (using a *dpp-LacZ* reporter, *dpp-Z*), the medium/high-HH target *ptc* and *engrailed* (*en*) which responds to the highest levels of HH in the anterior cells abutting the A/P boundary.

While expression of GFP-FU has no effect (Claret et al., 2007), coexpression of GFP-FU with T48 promotes the ectopic expression of *dpp-Z* and *ptc* throughout the whole anterior compartment (Figures 6A and 6B). However, it strongly reduces the anterior expression of *en* (Figure 6C) and accordingly leads to a stronger accumulation of CI-FL along the A/P border, indicating the loss of CI-A (Figure 6D). By contrast, trapping FU with NVR1 has no effect on *dppZ* and *ptc* expression and only weakly reduces the anterior *en* expression and CI-A (Figures 6E-6H).

**Figure 6:**
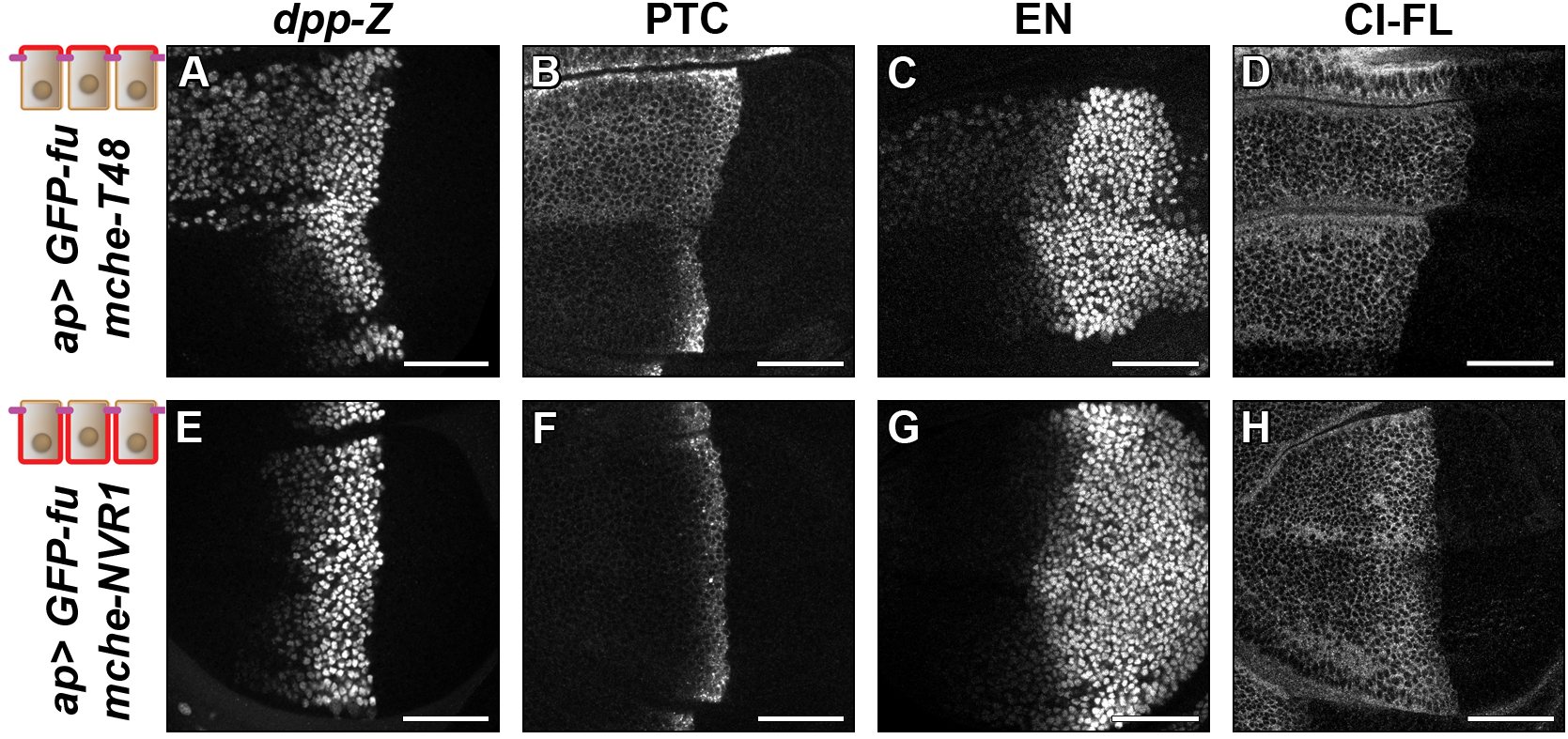
Trapping the FU kinase to the apical region is sufficient to promote low-medium but not high HH/SMO signaling. Confocal images of discs expressing GFP-fused with mche-T48 **(A-D)** or GFP-fu with mche-NVR1 **(E-H)** immunolabeled for the β-galactosidase (dpp-Z, to report dpp expression) (A and E), PTC (B and F), EN (C and G) and unprocessed CI allowing the detection of both CI-FL and CI-A, noted CI-FL here, D and H). The discs shown in (A) and (E) also expressed the *dpp-lacZ* reporter. The normal pattern of expression of these genes is visible in the ventral region of the discs. See also Figure S4.

In summary, anchoring FU to the apical membrane, but not to the basolateral one, increases both the levels of SMO and its localization at the cell surface. It also leads to its constitutive activity, promoting low-medium HH signaling but blocking very-high HH signaling. This suggests (i) that only apical FU can activate SMO and stabilize it at the cell surface and (ii) that both the high HH-induced basal enrichement of SMO and the expression of high HH targets may require a second input of FU on SMO in the basolateral region.

## DISCUSSION

Understanding how HH controls the fate of SMO in a polarized epithelium is central to understanding how this GPCR can be activated both in physiological and pathological conditions. Here we provide evidence that support a model (Figure 7) whereby (i) SMO is initially addressed to the apical membrane before being transcytosed to the basolateral region, (ii) HH controls the post-endocytic fate of SMO likely by enhancing its recycling, especially in the basolateral region, (iii) very high levels of HH favors a local trapping of SMO in the most basal region and finally that (iv) these effects rely on a SMO phosphorylation-barcode determined by the sequential action of the PKA/CKI and FU kinases, with FU acting in a two-step manner.

**Figure 7:**
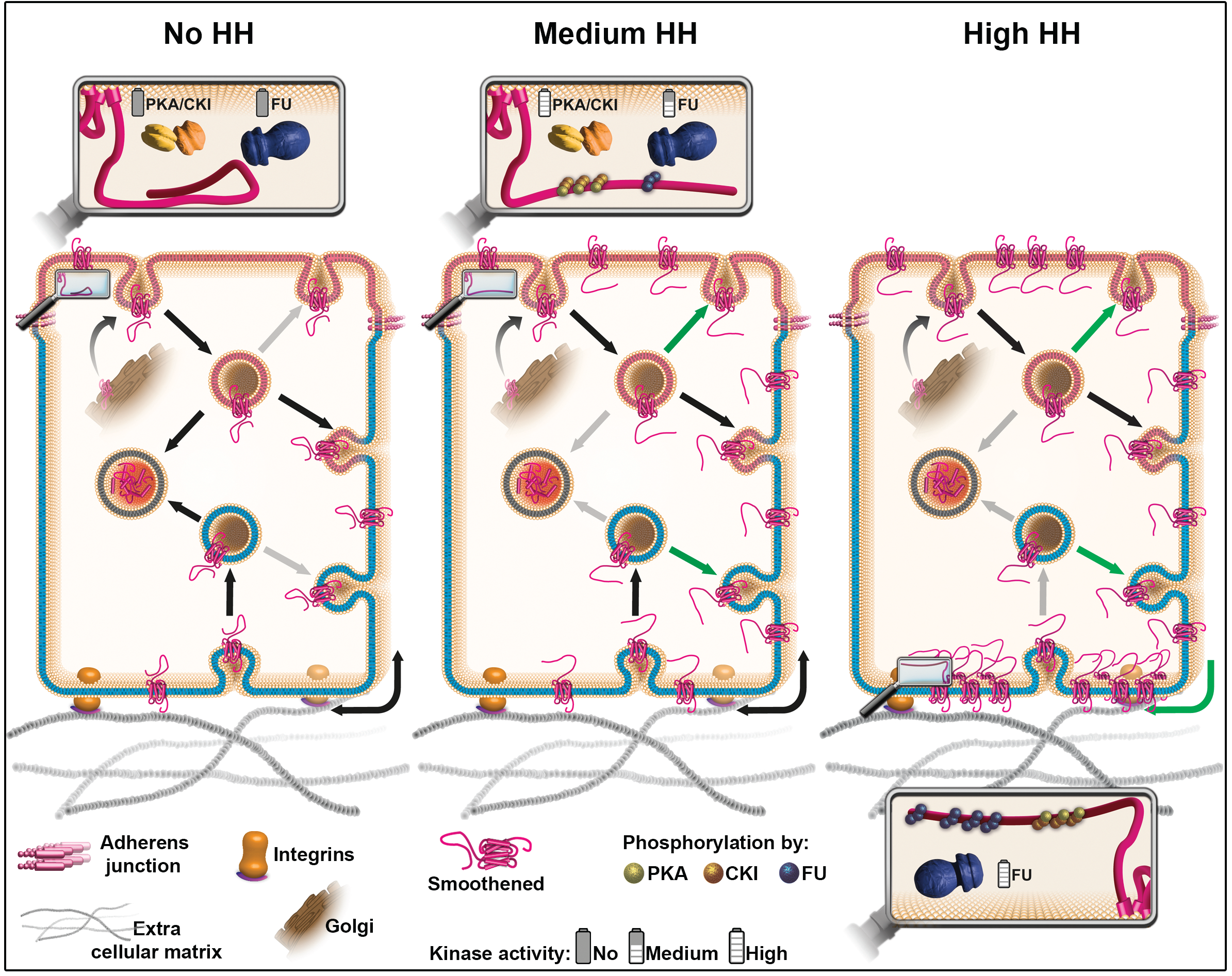
Model: HH controls the fate of endocytosed SMO via a multikinase phosphorylation barcode. SMO is initially targeted (curved arrow) from the Golgi compartment to the apical plasma membrane (presented in pink), where it is endocytosed (pink membrane vesicle). Endocytosed SMO can then be sent to the lysosome for degradation (grey membrane vesicle), recycled to the apical membrane or transcytosed to the basolateral membrane (represented in blue), where it can in turn, be re-endocytosed (blue membrane vesicle) to be recycled or degraded. Note that SMO can diffuse in the basolateral membrane (black double-headed arrow). HH controls several of these steps via the graded action of the PKA/CKI and FU kinases. In absence of HH (left panel), a large fraction of endocytosed SMO is targeted for degradation at the expense of its recycling (reduced recycling is indicated by grey arrows). In that situation, SMO cytotail is not phosphorylated and adopts a closed, inactive conformation. Oppositely, in presence of HH (see middle and right panels) the degradation of endocytosed SMO is reduced (grey arrows), in favor of its recycling (green arrows), especially in the basolateral region. These effects of HH lead to an increased accumulation of SMO at the cell surface and require the phosphorylation of the SMO’s cytotail by the PKA/CKI. This triggers a change in the conformation of SMO’s cytotail, associated with its activation. In presence of intermediate levels of HH (middle panel), SMO accumulation and/or activation reaches a threshold that in turn activates FU, which then retroacts on SMO by partially phosphorylating it, further enhancing the stabilizing and activating effects of the PKA/CKI kinases. The fact that FU is required for that effect in the apical region, suggests that the PKA/CKI also act in this domain. In the presence of high levels of HH (right panel), FU is further activated and promotes the enrichment of SMO in the most basal region by further phosphorylating its cytotail, which likely increases its clustering. This favors SMO’s trapping in the most basal region, which leads to high levels of HH signaling. The green arrow on the bottom right corner indicates a putative diffusion trapping mechanism, the grey bottom arrow indicates that recycling may be subsequently reduced.

The stabilization of SMO induced by HH could result from a reduction of its internalization or of its degradation after internalization. Our results indicate SMO endocytosis is little affected by HH (except for a possible slight reduction of its basal endocytosis) and that HH acts on endocytosed SMO, shifting its fate towards recycling rather than degradation. This involves the phosphorylation of SMO by the PKA/CKI, whose effects are further enhanced by a secondary action of FU. Notably, phosphorylation of the β-adrenergic receptor by the PKA has also been shown to increase its recycling to promote its resensitization (Gardner et al., 2004).

We have previously provided evidence for a double positive SMO-FU feedback loop behind “high HH” signaling: FU recruitment at the plasma membrane by SMO leads to a first level of FU activation, which in turn further activates SMO, which further increases FU activation. Here, tethering of FU to the apical membrane -but not to the basolateral one-is sufficient to ectopically promote both the stabilisation of SMO at the cell membrane and the activation of low/medium HH targets. However, in contrast to what is seen in presence of very high levels of HH or when SMO is fully hyperphosphorylated (SMO^PKA-SD FU-SD^), apical FU does not promote high levels of HH signaling and does not lead to a basal enrichment of SMO. Together, these data support the existence of a second effect of FU on SMO, which would occur in the basolateral region and lead to a basal accumulation of SMO, promoting very high HH signaling. Note that the aPKC was also reported to positively modulate SMO activity and to favor (directly or indirectly) its basolateral localization (Jiang et al., 2014). However, contrarily to the phosphorylation of SMO by FU, the phosphorylation by the aPKC does not seem to affect the “high HH”-dependent basal localization of SMO, nor “high HH” signaling, suggesting that it acts at another step.

Although the entire basolateral membrane is overall considered as a unique membrane domain in which proteins and lipids freely diffuse, many examples of membrane subregionalisation exist (for review see Trimble and Grinstein, 2015). Here, we provide evidence that hyperactivated SMO can be enriched in the most basal region. It could in part be due to a partial reduction of its basal endocytosis, but it likely also involves other mechanisms that where shown to restrain the localization of transmembrane proteins (for review see Trimble and Grinstein, 2015). For instance, it could involve an active oriented displacement of endocytic vesicles carrying SMO and FU directly to the basal domain by the kinesin COS2, which is known to be required for high HH signaling and to transport SMO and FU along microtubules in cultured fly cells (Farzan et al., 2008). Alternatively, diffusion trapping or partitioning phenomena (for review see Trimble and Grinstein, 2015) are also known to lead to local protein enrichment, for instance in axons (Ashby et al., 2006). Here, the changes in SMO electrostatic charges, conformation and/or clustering, which result from its hyperphosphorylation, could favor one or several of these processes (Shi et al., 2013; Zhao et al., 2007).

Regardless of the mechanism leading to this basal subpopulation of SMO, our data show that its presence is correlated to high levels of SMO activation and to the basal gradient of HH. Although we cannot exclude that this basal localization is a consequence rather than a cause of SMO activation, for instance a desensitization mechanism, our data along with published results strongly suggest that it is crucial to promote high HH signaling. We propose that in presence of high levels of HH, SMO could be trapped in basal specialized lipid microdomains that enhance signaling, similarly to what has been shown for the regulation of several GPCR by lipids rafts (for review see Villar et al., 2016). This possibility is supported by many reports showing that, both in flies and in mammals SMO responds to changes in its lipidic environment (for review see Radhakrishnan et al., 2020 and Zhang et al., 2021) with an emerging key role of acessible cholesterol (Kinnebrew et al., 2021). Notably, in Drosophila wing imaginal disc cells, SMO also was shown to relocalize in response to HH to cholesterol–rich raft lipid microdomains in the plasma membrane, where it forms higher-order structures (e.g. oligomers) that are required for high HH signaling, but the apico-basal localization of these raft was not adressed (Shi et al., 2013). Such microdomains could constitute signaling platforms acting on SMO structure (Zhao et al., 2021) enhancing the oligomerisation of SMO and of the HTC bound to it (Shi et al., 2011) and /or the interaction of SMO with specific HH signaling protein(s) (as seen with the EvC proteins in the cilia, see below (Dorn et al., 2012; Yang et al., 2012)).

Several lines of evidence point toward the conservation of a diffusion-trapping mechanism that would lead to the activation of SMO via its subcompartimentalisation in specific membrane domains of the plasma membrane. Indeed, in response to Sonic HH, the entry of SMO in the primary cilium of mammalian cells, which also depends on its phosphorylation (Chen and Jiang, 2013), was shown to involve the lateral diffusion of SMO from the cell body into the cilium membrane (Milenkovic et al., 2009) and is followed by its spatial restriction in a ciliary distinct compartment named the EvC zone (Dorn et al., 2012; Yang et al., 2012). This involves the phosphorylation-dependent interaction of SMO with two components of this zone, EvC and EvC2, both acting downstream of SMO to alleviate the negative effects exerted by Suppressor of Fused (SUFU). In that respect, it is worth noting, that in the Drosophila wing imaginal disc, the negative effects of SUFU need to be suppressed by FU for high “HH signaling” (Alves et al., 1998).

## MATERIALS AND METHODS

### Drosophila strains and genetics

*UAS SNAP-smo*^*WT*^ (Chr. III (Sanial et al., 2017)). The *UAS SNAP-smo*^*PKA-SD/PKA-SD FU-SA/PKA-SD FU-SD*^ flies were generated in this work by BestGene Inc. All *smo* transgenes were introduced into the same landing site (9738) on 3R (at 99F8) using the PhiC31 integration system to ensure that they are expressed at similar levels (Bateman et al., 2006). Further details are given in Supplementary Information. The genotypes of the strains used in the different figures are listed in the Supplemental Table1.

### Wing discs labeling

For total SNAP labeling, third instar larvae were dissected in Shields and Sang M3 Insect cell complete medium. They were then incubated for 20 min at 25°C with the SNAP-Cell TMR-Star (NEB) at a concentration of 2 μM in Shields and Sang M3 Insect cell complete medium. The larvae were fixed 20 min with PFA 4%, followed by three 10 min washes (PBS with 0.3% Triton) before immunolabeling or direct imaging. For surface labeling, dissected larvae were incubated for 10 min at 25°C with the SNAP-Surface Alexa Fluor 488, at 3.3 μM (NEB), fixed, washed before either incubation with SNAP-Cell TMR-Star at 2 μM (NEB) and SNAP-Surface Block at 13 μM (NEB) for 20 min at 25°C (for intracellular labeling) or immunolabeling.

Immunolabeling was done as in (Sanial et al., 2017). Primary antibodies: mouse anti PTC, 1:100 (Apa 1, from Developmental Studies Hybridoma Bank (DHSB)) (Martın et al., 2001) mouse anti EN, 1:50 (4D9, DHSB) (Patel et al., 1989); rabbit anti Fused, 1:100 (from P. Therond (Ruel et al., 2003); rabbit anti-β-Galactosidase, 1:100 (085597-CF, MP); mouse anti DLG 1:50 (4F3, DHSB) (Parnas et al., 2001); mouse anti SMO, 1:100 (20C6, from DSHB) (Lum et al., 2003) ; rat anti full length CI-FL, 1:5 (2A1 from R. Holmgren, Northwestern University, Evanston, IL, USA (Motzny and Holmgren, 1995)). Secondary antibodies: goat anti-mouse IgG Alexa Fluor™ Plus 555 highly cross-absorbed (Invitrogen, A32727); goat anti-rat IgG Alexa Fluor™ Plus 647 (Invitrogen, A21247); goat anti-rabbit IgG Alexa Fluor™ Plus 555 highly cross-absorbed (Invitrogen, A32732); goat anti-rabbit IgG Alexa Fluor™ Plus 488 highly cross-absorbed (Invitrogen, A32731); goat anti-rabbit IgG Alexa Fluor™ Plus 647 highly cross-absorbed (Invitrogen, A21245), chicken anti-mouse IgG cross adsorbed 488 (Invitrogen, A-21200), donkey anti-mouse IgG 649 (Jackson Immuno, 715-495-151) at a concentration of 1:200.

### Analysis of SMO trafficking

For the pulse-chase experiments, dissected wing imaginal discs were incubated for 10 min at 25°C with SNAP-Surface Alexa Fluor 546 (3.3 μM, from NEB), rinsed and incubated for 15 min with SNAP-Surface Block at 13 μM before fixation and immunolabeling. For the study of the endocytosis of SMO, *shi*^*ts*^*/Y; apGal4/+; UAS SNAP-smo*^*WT*^*/+* and *shi*^*+*^*/Y; apGal4/+; UAS SNAP-smo*^*WT*^*/+* wandering L3 larvae were incubated 30°C for 30 min, before being dissected. After dissection, the wing imaginal discs were incubated with SNAP-Surface Alexa Fluor 546 for 10 min at 30°C before fixation and immunolabeling. *apGal4, gal80*^*ts*^*/UAS yfp-rab5*^*CA*^; *UAS SNAP-smo*^*WT*^*/+* and apGal4, *gal80*^*ts*^*/+; UAS SNAP-smo*^*WT*^*/+* larvae were kept at 18°C (P Zeidler, et al., 2004) for 7-8 days. Vials were then moved to 29°C to induce the GAL4-dependent transcription of the UAS constructs for 24 h.

### Image acquisition, processing and quantification

XZ and YZ stacks of discs were acquired with sections every 1 μm using the confocal SP5 AOBS, 40x objective. Microscope settings were chosen to allow the highest fluorescence levels to be imaged under non-saturating conditions. Image data were processed and quantified using ImageJ software (National Institute of Health).

An ImageJ macro was designed to quantify SNAP-SMO fluorescence from the Z projections (average intensity) of the stack (8 sections). Firstly, the macro determines the shape and limits of the disc. It then asks the user to select a point in the disc (selected at the A/P boundary) from which rectangular regions from apical to basal, with the same width, will be drawn across the disc. Following this, each region is divided into three smaller ones: the apical/subapical, lateral and the basal subregions. The macro considers apical/subapical and basal subregions to be 15% and 10%, respectively, of the disc’s thickness. The 15% value was fixed based on DCAD (adherens junctions) and DLG (septate junctions) immunostaining. The 10% value is arbitrary and based on image analysis. The macro proceeds to measure the raw integrated density and the mean density of the different regions. Finally, with the help of CI immunostaining, four regions are selected across the disc (as shown in Figure 1H and S1A) to do the quantifications.

### Statistics and data representation

Statistical analysis was carried out using GraphPad Prism 8. Sample size was chosen large enough (n ≥ 8) to allow assessing statistical significance. Sample number and p-values are indicated in either the figure or the figure legend for each experiment. N-numbers indicate biological replicates, meaning the number of biological specimens evaluated (e.g. the number of wing discs). When comparing the anterior and posterior compartments within the same disc a Paired t test (for the Mean Intensities) or a Wilcoxon matched-pairs signed rank test (for the Relative Intensities) was used (Figure 1). When comparing different discs, a Mann-Whitney test was used (Figures 2, 3, 4). The p-values are shown in the Supplemental Table 2.

## ACKNOWLEDGMENTS

We are grateful to Drs. M. Crozatier, A. Guichet, D. Hipfner, J. Jian, P. Therond, F. Schweisguth for generously sharing their fly lines and reagents; to A. Benhmerah, A. Guichet and S. Léon and our colleagues from the Institut Jacques Monod for insightful discussions, Andréa Mialet and Severine Nozownik for their technical help. We are very grateful to Drs. G. D’Angelo and R. Holmgren for sharing their expertise and for their insightful advice. The Apa 1, 4D9, 4F3 and 20C6 monoclonal antibodies were obtained from the Developmental Studies Hybridoma Bank, created by the NICHD of the NIH and maintained at The University of Iowa, Department of Biology, Iowa City, IA 52242. Drosophila embryo injections were carried out by BestGene Inc. We acknowledge the ImagoSeine core facility of Institut Jacques Monod, member of France-BioImaging (ANR-10-INBS-04) and certified IBiSA.

## AUTHOR CONTRIBUTIONS

Conceptualization MGA, IB and AP; methodology MS and MGA; investigation MGA, MS, IB, SC and VC; formal analysis MGA, IB, AP and MS; writing, review and editing: IB, AP, MGA, MS; visualization MS and IB, supervision IB and AP; project administration and funding acquisition AP.

