## Supplemental information for "High Hedgehog signaling is transduced by a multikinase-dependent switch controling the apico-basal distribution of the GPCR Smoothened"

### Supplemental Experimental procedures

#### Drosophila strains and genetics

The following constructs and transgenic lines were previously published in (Sanial et al., 2017) or obtained from: *w*$^{1118}$ (Chr. I, (Hazelrigg et al., 1984)); *apGal4* (Chr. II, (Weihe et al., 2001)); *apGal4 gal80*$^{ts}$ (Chr. II, F. Schweisguth); *shibire*$^{ts}$ (van der Bliek and Meyerowrtz, 1991); *UAS YFP-rab5*$^{CA}$ (BL-9773); *UAS smurf RNAi* (BL-40905); *UAS mCherry-NVR1-GFP nanobody* (BL-68175) and *UAS mCherry-T48-GFP nanobody* (BL-68178) (Chr. III, (Harmansa et al., 2017); *UAS GFP-fused* (Chr. II, (Ruel et al., 2003)); *dpp-LacZ, apGal4* (Chr. II) was kindly given by R . Holmgren (*dpp-lac*Z is BL-12379). Complete genotypes are indicated in the supplemental Table 1.

#### Imaginal discs immunolabeling

Imaginal discs from third-instar wandering larvae were dissected in phosphate buffered saline (PBS), fixed 20 min at room temperature (RT) in 4% paraformaldehyde, washed three times 10 min in PBS + 0.3% Triton (PBST). They were incubated with the primary antibody overnight at 4°C. They were then washed three times for 10 min with PBST and incubated for 2 hr at RT with the secondary antibody in PBST before three washes 10 min in PBST and were mounted in Citifluor (Biovalley).

#### Plasmids

We used the Gateway Technology (Invitrogen following the manufacturer's instructions) to introduce the *SNAP-smo*$^{PKA\text{-}SD}$, *SNAP-smo*$^{PKA\text{-}SD\ FU\text{-}SA}$ or *SNAP-smo*$^{PKA\ PKA\text{-}SD\ FU\text{-}SD}$ transgenes in the vector *pUASt-GW-attB* (constructed by A. Brigui by insertion of the GW recombination cassette C3 at the EcoRI site of the pUASt-attB plasmid (GI EF362409)) for PhiC31 germline transformation, respectively. Prior to that, the PCR products obtained from the coding sequence (without the termination codon) of a *smo* wild-type cDNA were inserted into *pENTR/D-TOPO* by directional TOPO Cloning. Mutations leading to the S to A and S to D changes of the PKA/CKI sites were inserted into *pENTR/D-TOPO-snap-smo* by replacement of a region by the similar region coming from *smo*$^{PKA\text{-}SD/SA}$ from (Jia et al., 2004), leading to *pENTR/D-TOPO-snap-smo*$^{PKA\text{-}SA/PKA\text{-}SD}$. The mutations leading to the S to A and S to D replacements of the Fused phosphosites were introduced into *pENTR/D-TOPO-snap smo*$^{PKA\text{-}SD}$ by replacement of a region by the similar region coming from *pENTR/D-TOPO-smo smo*$^{FU\text{-}SD/SA}$ (Sanial et al., 2017) leading to *pENTR/D-TOPO-snap-smo*$^{PKA\text{-}SD\ FU\text{-}SA}$ *and pENTR/D-TOPO-snap-smo*$^{PKA\text{-}SD\ FU\text{-}SD}$. All constructs were checked by sequencing the fragments produced by PCR and their junctions.

**Fiji quantification macro**

```
nbBand=9;
pourcentApical=0.15;
pourcentBasal=0.10;

run("Set Measurements...", "  redirect=None decimal=9");

myImage=getTitle();
selectWindow(myImage);
run("Split Channels");
close();
close();

segmentation("C1-"+myImage);
nbBand = createROI2("C1-"+myImage,"Mask of imgForThreshold",nbBand);
run("Z Project...", "projection=[Average Intensity]");
for(n=0;n<nbBand;n++){
	setResult("image",n,myImage);
	ApicalBasal("AVG_C1-"+myImage,n,pourcentApical,pourcentBasal);
}
for(n=0;n<nbBand;n++){
	roiManager("Select",0);
	roiManager("Delete");
}
intensityMap("AVG_C1-"+myImage);



function segmentation(nameImage){
```

```
        selectWindow(nameImage);
        run("Z Project...", "projection=[Average Intensity]");
        selectWindow("AVG_"+nameImage);
        run("Duplicate...", "duplicate channels=1");
        rename("imgForThreshold");
        run("Median...", "radius=20 stack");
        run("8-bit");
        run("Auto Local Threshold", "method=Sauvola radius=128 parameter_1=0.25 parameter_2=256 white");
        run("Analyze Particles...", "size=300000-Infinity pixel show=Masks");
        run("Invert LUT");
        run("Fill Holes");
        run("Options...", "iterations=100 count=2 black pad do=Close");
        selectWindow("imgForThreshold");close();
    }



    function createROI2(nameImage,nameMask,nbBand){
        selectWindow(nameMask);
        run("Create Selection");
        selectWindow(nameImage);
        run("Restore Selection");
        roiManager("Add");
        roiManager("Select", 0);
        roiManager("Rename", "ROITissus");
        roiManager("Deselect");run("Select None");
        setTool("point");
        waitForUser("Wait on click", "Please click on the middle of a section and click on OK.");
        getSelectionCoordinates(xclick, yclick);run("Select None");
        print("Point du clic : "+xclick[0]);
        print("click="+xclick[0]);

        width=1024/nbBand;
        print("Width="+width);
        shift=(xclick[0]%width)-(width/2);
        print("shift="+shift);
        roiManager("Select", 0);
        width=1024/nbBand;
        if(shift>=0){
            run("Specify...", "width="+width+" height=1024 x="+shift+" y=0 slice=1");
        }
        else{
            run("Specify...", "width="+width+" height=1024 x="+(width+shift)+" y=0 slice=1");
            print("x=x="+(width));
```

```
    }
    roiManager("Add");
    if(shift!=0){nbBand=nbBand-1;}
    for(i=1;i<=nbBand;i++){
        if(shift>=0){
            x=width*(i-1)+shift;    }
        else{
            x=width*(i-1)+width+shift;}
        print("xx="+x);
        roiManager("Select", 1);
        run("Specify...", "width="+width+" height=1024 x="+x+" y=0 slice=1");
        roiManager("Update");
        roiManager("Rename", "Largeband"+i);
        roiManager("Select", newArray(0,1));
        roiManager("AND");
        run("To Bounding Box");
        roiManager("Add");
        roiManager("Deselect");
        roiManager("Select", i+1);
        roiManager("Rename", "Band"+i);
        roiManager("Deselect");
    }
    roiManager("Select", 0);roiManager("Delete");
    roiManager("Select", 0);roiManager("Delete");
    roiManager("Show All with labels");
    return nbBand;
}



function ApicalBasal(nomImage,i,pourcentApical,pourcentBasal){
    setResult("Band", i, i+1);////
    selectWindow(nomImage);
    roiManager("Show None")
    roiManager("Select", i);
    getRawStatistics(nPixels, mean, min, max, std, histogram);
    setResult("sum Band", i, nPixels*mean);////
    setResult("mean Band", i, mean);////
    Roi.getBounds(x, y, width, height);
    makeRectangle(x, y, width, round(height*pourcentApical));
    getRawStatistics(nPixels, mean, min, max, std, histogram);
    setResult("mean Apical", i, mean);////
    setResult("sum Apical", i, nPixels*mean);////
    roiManager("Add");    roiManager("Select", roiManager("count")-1);
    roiManager("Rename", "Band"+i+1+"_Apical");
    roiManager("Select", i);
```

```
        makeRectangle(x, y+round(height*(1-pourcentBasal)), width,
round(height*pourcentBasal));
        getRawStatistics(nPixels, mean, min, max, std, histogram);
        setResult("mean Basal", i, mean);////
        setResult("sum Basal", i, nPixels*mean);////
        roiManager("Add");roiManager("Select", roiManager("count")-1);
        roiManager("Rename", "Band"+i+1+"_Basal");

        roiManager("Select", i);
        makeRectangle(x, y+round(height*pourcentApical), width, round(height*(1-
(pourcentApical+pourcentBasal))));
        getRawStatistics(nPixels, mean, min, max, std, histogram);
        setResult("mean Medial", i, mean);////
        setResult("sum Medial", i, nPixels*mean);////
        roiManager("Add");roiManager("Select", roiManager("count")-1);
        roiManager("Rename", "Band"+i+1+"_Medial");

        updateResults();
        roiManager("Deselect");
    }



    function intensityMap(nomImage){

        selectWindow(nomImage);

        getDimensions(width, height, channels, slices, frames);

        newImage("IntensityMap", "16-bit black", width, height, 1);


        for(i=0;i<roiManager("count");i++){

                selectWindow(nomImage);

                roiManager("Select", i);

                getStatistics(area, mean, min, max, std, histogram);

                selectWindow("IntensityMap");

                roiManager("Select", i);
```

```
            setColor(mean);fill();
        }
        roiManager("Deselect");
        run("Select None");
        run("Fire");
        run("Enhance Contrast", "saturated=0.35");
    }
```

### Legends of supplemental Figures

**Figure S1: Apico-basal distribution of SMO in the wing imaginal disc epithelium.**

**(A)** Definition and representation of the different regions of the wing imaginal disc studied here. Upper panel: HH produced by the cells from the posterior compartment diffuses into the anterior region where it controls CI fate in a dose-dependent manner (the abbreviations and color code are indicated in Figure 1). Middle panel: Z section of a wing imaginal disc labeled for SNAP-SMO and showing the four regions used for the quantifications. Lower panel: definition of the different regions along the the apico-basal axis of the wing imaginal disc epithelium. The apical region corresponds to 15 % of the epithelium height (based on DLG staining), the basal region was arbitrarily defined as the 10% most basal region and the lateral region as (75% of the disc height) in between.

**(B-C)** Apico-basal distribution of endogenous SMO. Confocal images (XZ section) of wing imaginal discs immunolabeled for endogenous SMO (B, green in the merge image B") and CI-FL (B', blue in merge image B"), as indicated. Mean (C) and relative (C') intensities of endogenous SMO along the apico-basal axis in the far anterior and P regions. Note that, contrarily to *SNAP-smo* driven by *apGal4*, endogenous *smo* is expressed in the peripodial membrane (arrow).

**(D-D")** Mean intensity of Total (D), Intra (D') and Surf (D") SNAP-SMO along the apico-basal axis in the FA and P regions of the disc. The discs used for this quantification correspond to the same discs used in the graphs of Figures 1 E to 1E".

**(E)** Mean intensity of Surf SNAP-SMO along the apico-basal axis in the R, F, A and P regions of the wing imaginal disc corresponds to the same discs than in the graph of Figure 1H.

**(F)** Schematic representation of the method used for sequential labeling of Surf and Intra SNAP-SMO. Note that Surf SNAP-SMO corresponds to the SNAP-SMO molecules that were at least transiently present at the plasma membrane during the labeling time.

**Figure S2: Effect of *shi*$^{ts}$ and *rab5*$^{CA}$ on Surf SNAP-SMO$^{WT}$**

**(A-B)** False color representation of XY confocal images of wing imaginal disc from *shi*$^{+}$*; apGal4; UAS SNAP-smo*$^{WT}$ (A) or *shi*$^{ts}$*; apGal4; UAS SNAP-smo*$^{WT}$ (B) male flies put at restrictive temperature before dissection and labeling of Surf SNAP-SMO.

**(C-D)** XY confocal images of wing imaginal discs from *apGal4, Gal80ts; SNAP-smo*$^{WT}$ (C) or *apGal4, Gal80ts/UAS YFP-rab5*$^{CA}$*; UAS SNAP-smo*$^{WT}$ (D-D”) flies put at restrictive temperature before dissection and Surf SNAP-SMO labeling.

**(E)** Quantification of the apico-basal distribution of Surf SNAP-SMO in wing imaginal disc of *shi*$^{+}$*; apGal4; UAS SNAP-smo*$^{WT}$ (ctr) or *shi*$^{ts}$*; apGal4; UAS SNAP-smo*$^{WT}$ kept at 18°C. The same distribution is seen for the two genotypes, indicating that the effects in Figures 2B, 2C and 2C’ are due to the inactivation of the *shi*$^{ts}$ allele. N=8 for the controls and n=7 for the *shi*$^{ts}$ flies.

**Figure S3: False color representation of the accumulation at the cell surface of wild-type and phosphomutants forms of SNAP-SMO** (as indicated).

**(A-D)** XY section.

**(A’-D’)** XZ projection.

All discs were imaged in the same conditions.

**Figure S4: FU and SMO colocalization and effect on the subcellular localization of SMO of trapping the FU kinase to the apical or basolateral region**

**(A-A’)** Confocal images of wild-type discs immunolabeled for FU. XY section in (A) and XZ section in (A’).

**(B-B”)** XY confocal images of SMO (left panels) and FU (center panels) in wing imaginal discs of wild-type flies. Merge images are shown in the right panels. Sections at different levels of the apico-basal axis are shown (apical B, medial B’ and basal B”). Imaging was done using the LSM980 spectral Airyscan 2, 63X. Scale bar: 10 µm.

**(C-C”, D-D”)** YZ confocal images of wing imaginal discs coexpressing (using the apGal4 driver) *GFP-fu* with *mCherry-T48 (C-C”’)* or *mCherry-NVR1 (D-D”’)*. The Z sections correspond to dorso-ventral sections in the posterior compartment with the apical left, and dorsal up. mCherry is shown in (C and D, red in the merge image C” and D”), GFP-FU in (C’ and D’, green in the merge images C” and D”) and immunolabeled endogenous SMO in (C”’ and D”’).

**(E, E’**) Quantification of the mean intensity in the far anterior and posterior regions and of the relative intensity (E”) in the far anterior region. Note that the quantification method excluded the endogenous SMO labeling in the peripodial membrane.

### Supplemental Tables

**Supplemental Table 1: Fly genotypes**

| | |
|---|---|
| **Figure 1** | |
| | *apGal4/+; UAS SNAP-smo*$^{WT}$*/+* |
| **Figure 2** | |
| 2A and C | *shi*$^{+}$*/Y; apGal4/+; UAS SNAP-smo*$^{WT}$*/+* |
| 2B and C | *shi*$^{ts}$*/Y; apGal4/+; UAS SNAP-smo*$^{WT}$*/+* |
| 2D and F | *apGal4, Gal80*$^{ts}$*/+; SNAP-smo*$^{WT}$*/+* |
| 2E-E" and F | *apGal4, Gal80*$^{ts}$*/UAS YFP-rab5*$^{CA}$*; UAS SNAP-smo*$^{WT}$*/+* |
| 2G | *apGal4/+; SNAP-smo*$^{WT}$*/+* and<br>*apGal4/UAS smurf RNAi; UAS SNAP- smo*$^{WT}$ */+* |
| **Figure 3** | |
| | *apGal4/+; UAS SNAP-smo*$^{WT}$*/+* |
| **Figure 4** | |
| 4A and E-E" | *apGal4/+; UAS SNAP-smo*$^{WT}$*/+* |
| 4B and E-E" | *apGal4/+; UAS SNAP-smo*$^{PKA\text{-}SD}$*/+* |
| 4C and E-E" | *apGal4/+; UAS SNAP-smo*$^{PKA\text{-}SD\ FU\text{-}SA}$*/+* |
| 4D and E-E" | *apGal4/+; UAS SNAP-smo*$^{PKA\text{-}SD\ FU\text{-}SD}$*/+* |
| **Figure 5** | |
| 5A-A"', B-B"' | *apGal4/UAS GFP-fused; UAS mCherry-T48/+* |
| 5C-C"', C-C"' | *apGal4/UAS GFP-fused; UAS mCherry-NVR1/+* |
| **Figure 6** | |
| 6A | *dpp-Z, apGal4/UAS GFP-fused; UAS mCherry-T48/+* |
| 6B-D | *apGal4/UAS GFP-fused; UAS mCherry-T48/+* |
| 6E | *dpp-Z, apGal4/UAS GFP-fused; UAS mCherry-NVR1/+* |
| 6F-H | *apGal4/UAS GFP-fused; UAS mCherry-NVR1/+* |

**Supplemental Table 2: Statistical analysis**

| | | | | |
|---|---|---|---|---|
| **Figure 1** | | | | |
| E | | **** | <0.0001 | Paired t test |
| E' | | * | 0,0391 | Wilcoxon matched-pairs signed rank test |
| E'' | | *** | 0,0005 | |
| H | | **** | <0.0001 | |
| | | ** | 0,0016 | |
| | | *** | 0,0004 | |
| **Figure 2** | | | | |
| C | NO HH | ** | 0,0039 | Mann-Whitney test |
| | | **** | <0.0001 | |
| | | * | 0,0459 | |
| | | ** | 0,032 | |
| | HH | ** | 0,0014 | |
| | | **** | <0.0001 | |
| | | * | 0,0151 | |
| | | ** | 0,0093 | |
| C' | NO HH | **** | <0.0001 | |
| | HH | ** | 0,0022 | |
| F | | **** | <0.0001 | |
| | | *** | 0,0003 | |
| G | NO HH | **** | <0.0001 | |
| | | *** | 0,0002 | |
| | HH | **** | <0.0001 | |
| | | *** | 0,0008 | |
| | | *** | 0,0002 | |
| **Figure 3** | | | | |
| C | | **** | <0.0001 | Mann-Whitney test |
| D | NO HH | *** | 0,0005 | |
| | HH | **** | <0.0001 | |
| | | ** | 0,0085 | |
| | | *** | 0,001 | |
| **Figure 4** | | | | |
| E | Apical | **** | <0.0001 | Mann-Whitney test |
| | | ** | 0,0025 | |
| E' | lateral | ** | 0,0033 | |
| | | **** | <0.0001 | |
| E'' | Basal | *** | 0,0002 | |
| | | **** | <0.0001 | |
| For the comparison between WT -HH with WT +HH, the Wilcoxon test was used | | | | |

Figure S1:

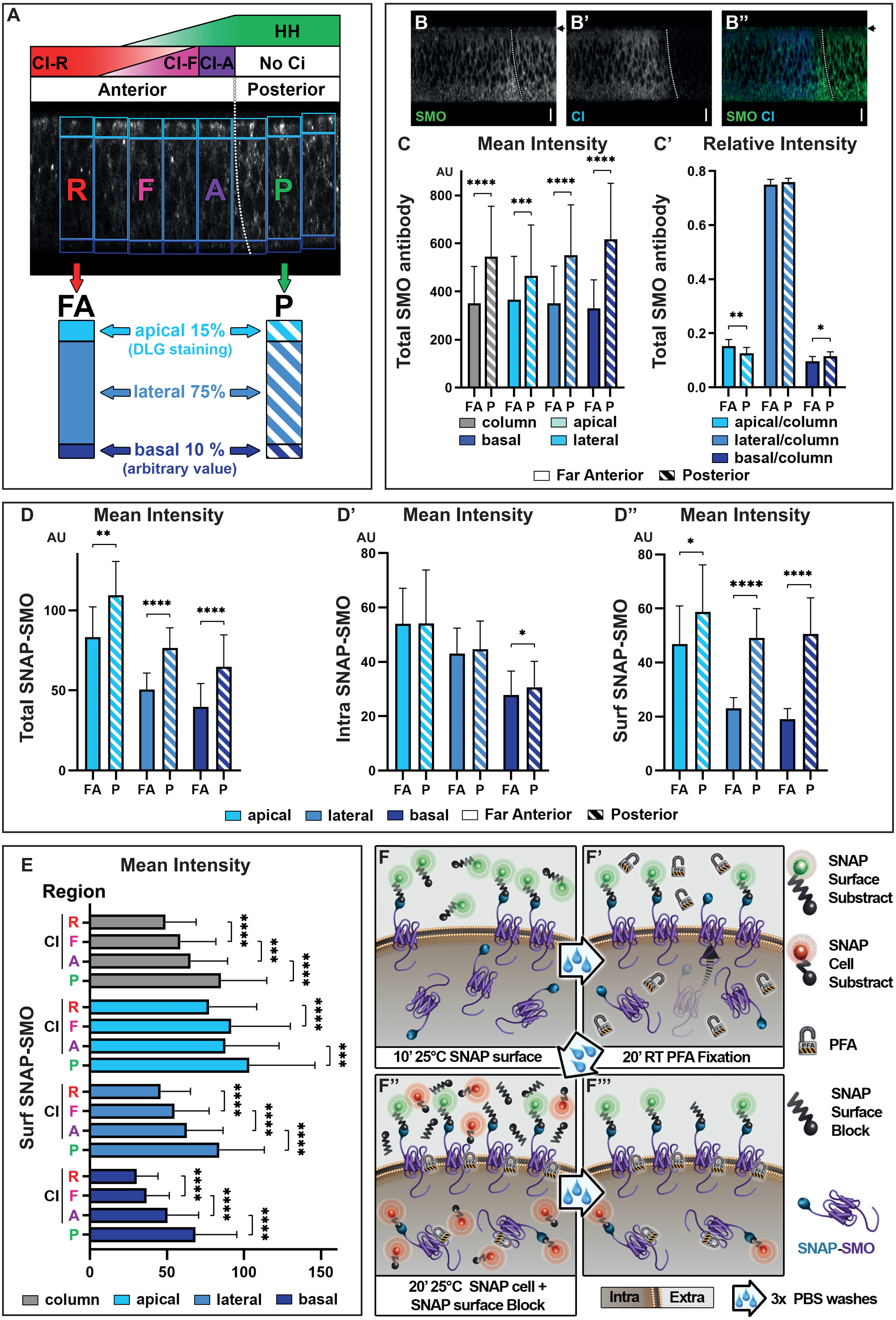

A
HH
CI-R
CI-F
CI-A
No Ci
Anterior
Posterior
R
F
A
P
FA
P
apical 15%
(DLG staining)
lateral 75%
basal 10 %
(arbitrary value)
B
B'
B''
SMO
CI
SMO CI
C
Mean Intensity
C'
Relative Intensity
Total SMO antibody
column
apical
basal
lateral
apical/column
lateral/column
basal/column
Far Anterior
Posterior
D
Mean Intensity
D'
Mean Intensity
D''
Mean Intensity
Total SNAP-SMO
Intra SNAP-SMO
Surf SNAP-SMO
apical
lateral
basal
Far Anterior
Posterior
E
Mean Intensity
Region
CI
Surf SNAP-SMO
column
apical
lateral
basal
F
F'
F''
F'''
10' 25°C SNAP surface
20' RT PFA Fixation
20' 25°C SNAP cell +
SNAP surface Block
SNAP Surface Substract
SNAP Cell Substract
PFA
SNAP Surface Block
SNAP-SMO
Intra
Extra
3x PBS washes

Figure S2:

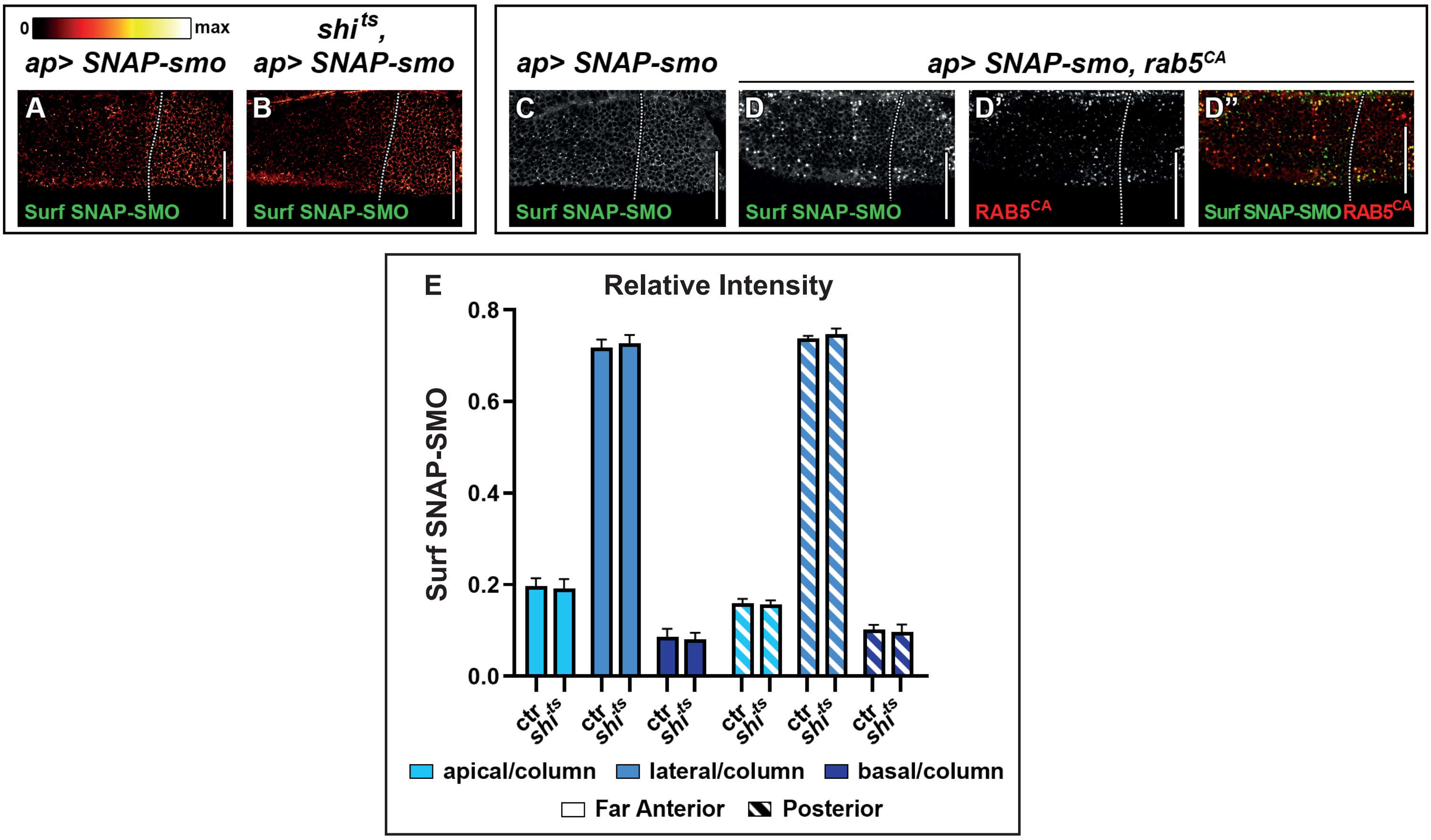

0
max
ap> SNAP-smo
shi ts,
ap> SNAP-smo
A
Surf SNAP-SMO
B
Surf SNAP-SMO
ap> SNAP-smo
ap> SNAP-smo, rab5CA
C
Surf SNAP-SMO
D
Surf SNAP-SMO
D'
RAB5CA
D''
Surf SNAP-SMO RAB5CA
E
Relative Intensity
Surf SNAP-SMO
0.0
0.2
0.4
0.6
0.8
ctr
shi ts
apical/column
lateral/column
basal/column
Far Anterior
Posterior

**Figure S3:**

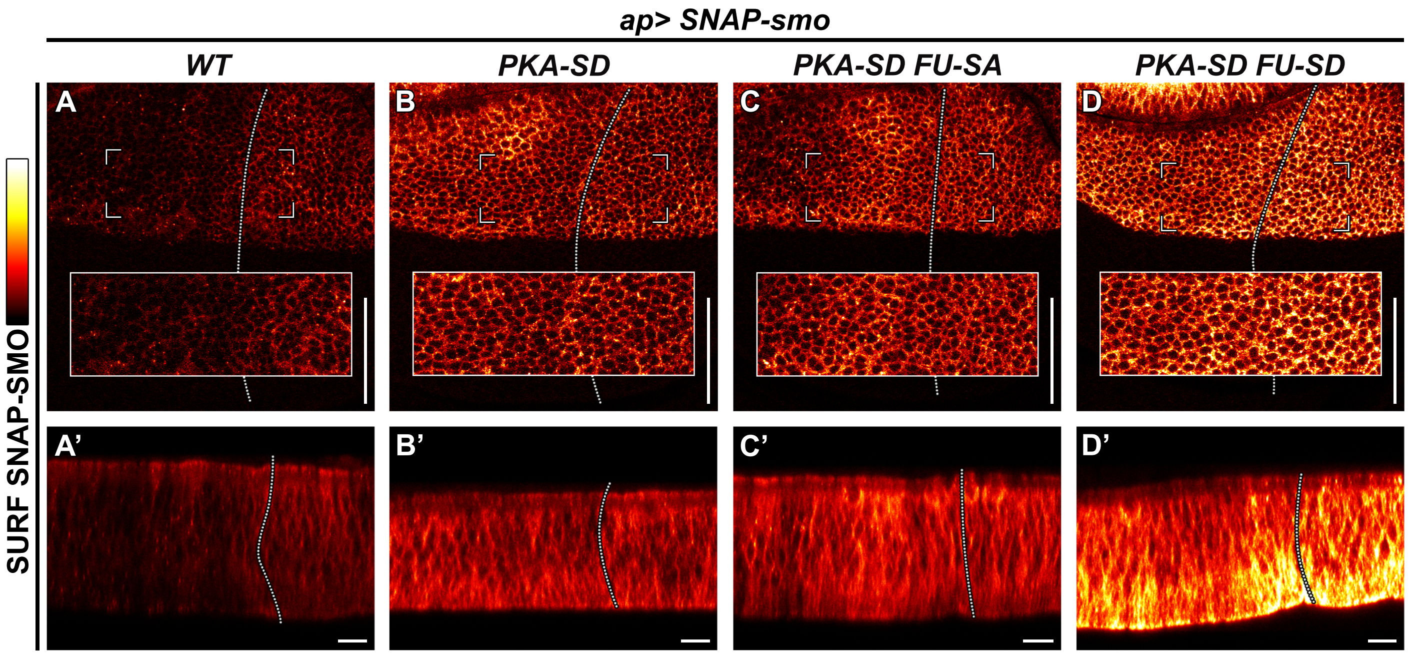

ap> SNAP-smo
WT
PKA-SD
PKA-SD FU-SA
PKA-SD FU-SD
A
B
C
D
A'
B'
C'
D'
SURF SNAP-SMO
